# Altered basal lipid metabolism underlies the functional impairment of naive CD8^+^ T cells in elderly humans

**DOI:** 10.1101/2020.09.24.311704

**Authors:** Francesco Nicoli, Mariela P. Cabral-Piccin, Laura Papagno, Eleonora Gallerani, Victor Folcher, Marion Dubois, Emmanuel Clave, Hélène Vallet, Justin J. Frere, Emma Gostick, Sian Llewellyn-Lacey, David A. Price, Antoine Toubert, Jacques Boddaert, Antonella Caputo, Riccardo Gavioli, Victor Appay

## Abstract

**Background:** Aging is associated with functional deficits in the naive T cell compartment, which compromise the generation of *de novo* immune responses against previously unencountered antigens. The mechanisms that underlie this phenomenon have nonetheless remained unclear.

**Methods:** Biochemical and functional properties of naive CD8^+^ T cells were characterized and compared between middle aged and older individuals.

**Findings:** We identified an age-related link between altered basal lipid metabolism in naive CD8^+^ T cells and their impaired responsiveness to stimulation, characterized by low proliferative potential and susceptibility to apoptosis. Reversal of the bioenergetic anomalies with lipid-altering drugs, such as rosiglitazone, improved the functional capabilities of naive CD8^+^ T cells in elderly subjects.

**Interpretation:** Interventions that favor lipid catabolism may find utility as adjunctive therapies in the elderly to promote vaccine-induced immunity against emerging pathogens or tumors.

**Funding:** A full list of the funding sources is detailed in the Acknowledgment section of the manuscript.

**RESEARCH IN CONTEXT:** *Evidence before this study:* Old subjects are highly susceptible to infections and tumors and usually present with low responses to vaccine. This is mainly due to the age-related loss of primary immune resources, i.e. a quantitative decline of naive CD8^+^ T cells. Nonetheless, few studies have also underlined, within this cell subset, qualitative defects in elderly subjects.

*Added value of this study:* Considering the well-demonstrated link between nutrient usage and lymphocyte functions, we characterized the bioenergetics features of old naïve CD8^+^ T cells. Our data show an age-dependent altered basal metabolism in this cell subset, mostly at the levels of fatty acids and mitochondrial functions. These alterations were associated with functional defects which were partially reverted through the use of lipid-lowering strategies.

*Implications of all the available evidence:* This study highlights the potential role of an altered cellular lipid metabolism in immunosenescence, providing clues to understand the epidemiological profile of emerging infections or tumors and to develop preventive and therapeutic strategies based on metabolic manipulation.

## INTRODUCTION

Life expectancy has increased considerably over the last century as a consequence of advances in medicine and improved public health systems. However, old age is associated with a high prevalence of chronic diseases and an increased susceptibility to cancer and emerging pathogens, such as SARS-CoV-2 [1]. Age-related deficits in the immune system are thought to play a key role in the development of many pathological conditions [2–4]. Immune aging is characterized by a progressive erosion of the naive CD8^+^ T cell compartment, which impairs *de novo* immune responses against newly encountered antigens [5,6]. In addition to a decline in absolute numbers [7], naive CD8^+^ T cells in elderly individuals exhibit impaired differentiation in response to T cell receptor (TCR)-mediated activation [5].

A growing body of evidence indicates that lymphocyte metabolism is a key determinant of immune functionality [8–11]. Systemic metabolic disturbances are common in elderly individuals, and increased levels of adipokines and proinflammatory lipid species in particular have been implicated as critical mediators of inflammaging, which is thought to exacerbate many age-related diseases [12]. In this study, we investigated the bioenergetic features of naive CD8^+^ T cells in middle-aged and elderly humans, aiming to establish a link between metabolic disturbances and age-related functional impairments. Naive CD8^+^ T cells displayed specific metabolic abnormalities in elderly people, in particular enhanced lipid influx and storage, accompanied by reduced proliferation and increased susceptibility to apoptosis upon activation. Importantly, these deficits were mitigated in the presence of lipid-altering drugs, opening potential therapeutic avenues to slow the process of immunosenescence.

## METHODS

### Human subjects and samples

Two groups of healthy volunteers were enrolled in this study: (i) middle-aged Caucasians (19 to 55 years old; mediane: 39); and (ii) elderly Caucasians (65 to 95 years old; mediane: 82). Individuals with malignancies, acute diseases, or severe chronic diseases, such as atherosclerosis, congestive heart failure, poorly controlled diabetes mellitus, renal or hepatic disease, various inflammatory conditions, or chronic obstructive pulmonary disease, as well as individuals on immunosuppressive therapy, were excluded from the study. PBMCs were isolated from venous blood samples via density gradient centrifugation according to standard protocols and cryopreserved in complete medium supplemented with dimethyl sulfoxide (DMSO; 10% v/v; Sigma-Aldrich) and fetal calf serum (FCS; 20% v/v; Sigma-Aldrich). Complete medium (R+) consisted of RPMI 1640 supplemented with non-essential amino acids (1% v/v), penicillin/streptomvcni (100 U/mL), L-glutamine (2 mM), and sodium pyruvate (1 mM) (all from Thermo Fisher Scientific).

### Flow cytometry and cell sorting

PBMCs were stained for surface markers using combinations of the following directly conjugated monoclonal antibodies: anti-CCR7–BV650 (clone 3D12; BD Biosciences), anti-CCR7–PE-Cy7 (clone 3D12; BD Biosciences), anti-CD3–BV605 (clone SK7; BD Biosciences), anti-CD8–APC (clone RPA-T8; BD Biosciences), anti-CD8–APC-Cy7 (clone SK1; BD Biosciences), anti-CD8–FITC (clone RPA-T8; BD Biosciences), anti-CD27–AF700 (clone O323; BioLegend), anti-CD27-PE (clone M-T271; BD Biosciences), anti-CD45RA–ECD (clone 2H4LDH11LDB9; Beckman Coulter), anti-CD45RA–PerCP-Cy5.5 (clone HI100; eBioscience), anti-CD45RA–V450 (clone HI100; BD Biosciences), anti-CD49b–PE-Cy7 (clone 9F10; BioLegend), anti-CD57–Pacific Blue (clone HCD57; BioLegend), and anti-CD95–FITC (clone DX2; BD Biosciences). Naive CD8^+^ T cells were defined as CD3^+^ CD8^+^ CD27^+^ CD45RA^+^ CCR7^+^ in most experiments and further identified as CD49b^-^ CD57^-^ CD95^-^ for gene expression studies and intracellular measurements of T-bet. Non-viable cells were eliminated from the analysis using LIVE/DEAD Fixable Aqua (Thermo Fisher Scientific). Intracellular stains were performed using anti-T-beteFluor66O (clone 4B10; eBioscience) in conjunction with a Transcription Factor Buffer Set (BD Biosciences). Samples were acquired using an LSR Fortessa or a FACSCanto II (BD Biosciences). Naive CD8^+^ T-cells were flow-sorted using a FACSAria II (BD Biosciences). Data were analyzed using FACSDiva software version 7 (BD Biosciences) and/or FlowJo software version 10 (FlowJo LLC).

### Proliferation assays

PBMCs were labeled with Cell Proliferation Dye (CPD) eFluor450 (Thermo Fisher Scientific) and stimulated for 4 days with plate-bound anti-CD3 (clone OKT3; Thermo Fisher Scientific) in the absence or presence of CTAB (1 μM; Sigma-Aldrich). In some experiments, cells were precultured in AIM-V medium (Thermo Fisher Scientific) supplemented with bovine serum albumin (BSA; 10% v/v; Sigma-Aldrich) for 1 day in the absence or presence of palmitic acid (300 μM; Sigma-Aldrich), and in other experiments, cells were precultured in AIM-V medium (Thermo Fisher Scientific) without BSA supplementation for 2 days in the absence or presence of rosiglitazone (40 μM; Sigma-Aldrich). Proliferation was measured using flow cytometry to quantify the dilution of CPD.

### Activation assays

PBMCs were stimulated for 24 hr with plate-bound anti-CD3 (clone OKT3; Thermo Fisher Scientific) in the absence or presence of fenofibrate (50 μM; Sigma-Aldrich) or rosiglitazone (40 μM; Sigma-Aldrich). The expression of the standard activation markers CD69 and CD134 (OX40) was quantified on the cell surface using anti-CD69–FITC (clone L78; BD Biosciences) and anti-CD134–BV711 (clone ACT35; BD Biosciences). Intracellular staining for activated caspase-3, used as marker of susceptibility to apoptosis upon activation, was performed using anti-active caspase-3–PE (clone C92-605; BD Biosciences) in conjunction with a Cytofix/Cytoperm Fixation/Permeabilization Solution Kit (BD Biosciences).

### Metabolism assays

To determine glucose uptake, PBMCs were incubated for 20 min at 37°C in phosphate-buffered saline (PBS) containing 2’-(N-(7-nitrobenz-2-oxa-1,3-diazol-4-yl)amino)-2-deoxyglucose (2-NBDG; 50 μM; Thermo Fisher Scientific). To determine FA uptake, PBMCs were incubated for 20 min at 37°C in PBS containing 4,4-difluoro-5,7-dimethyl-4-bora-3a,4a-diaza-*s*-indacene-3-hexadecanoic acid (BODIPY FL C16; 1 μM; Thermo Fisher Scientific). To determine neutral lipid content, PBMCs were incubated for 20 min at 37°C in PBS containing 4,4-difluoro-1,3,5,7,8-pentamethyl-4-bora-3a,4a-diaza-*s*-indacene (BODIPY 493/503; 10 μM; Thermo Fisher Scientific). To determine mitochondrial mass, PBMCs were incubated for 30 min at 37°C in R+ containing Mitotracker Deep Red (500 nM; Thermo Fisher Scientific). To determine mitochondrial membrane potential, PBMCs were incubated for 30 min at 37°C in R+ containing tetramethylrhodamine, methyl ester, perchlorate (TMRM; 25 nM; Thermo Fisher Scientific). To determine mTOR activity, PBMCs were incubated for 10 min at 37°C in Cytofix Fixation Buffer (BD Biosciences), washed, incubated for 30 min at 4°C in Phosflow Perm Buffer III (BD Biosciences), washed again, and stained for 1 hr at room temperature with anti-pS6–Pacific Blue (clone D57.2.2E; Cell Signaling Technology).

### Peptides and tetramers

All peptides were synthesized at >95% purity (BioSynthesis Inc.). The EV20 peptide (YTAAEELAGIGILTVILGVL, Melan-A_21–40/A27L_) was used for *in vitro* priming studies. Fluorochrome-labeled tetrameric complexes of HLA-A*02:01–EV10 (ELAGIGILTV, Melan-A_26–35/A27L_) were generated in-house as described previously [13].

### *In vitro* priming of antigen-specific CD8^+^ T cells

Naive precursors specific for HLA-A2–EV10 were primed *in vitro* using an accelerated dendritic cell (DC) coculture protocol as described previously [9,14,15]. Briefly, thawed PBMCs were resuspended at 5 × 10^6^ cells/well in AIM-V medium (Thermo Fisher Scientific) supplemented with Flt3 ligand (Flt3L; 50 ng/ml; R&D Systems) in the absence or presence of rosiglitazone (40 μM; Sigma-Aldrich) or IL-7 (20 ng/mL; R&D Systems). After 24 hr (day 1), the Melan-A peptide EV20 (1 μM) was added to the cultures, and DC maturation was induced using a standard cocktail of inflammatory cytokines, incorporating IL-1β (10 ng/mL), IL-7 (0.5 ng/mL), prostaglandin E2 (PGE2; 1 μM), and TNF (1,000 U/mL) (all from R&D Systems), or ssRNA40 (TLR8L; 0.5 μg/mL; InvivoGen). The cultures were supplemented on day 2 with FCS (10% v/v; Sigma-Aldrich). Medium was replaced every 3 days thereafter with fresh R+ containing FCS (10% v/v; Sigma-Aldrich). Antigen-specific CD8^+^ T cells were characterized via flow cytometry on day 10.

### RNA extraction and qPCR analysis

PBMCs were activated for 5 hr with plate-bound anti-CD3 (clone OKT3; Thermo Fisher Scientific). RNA was extracted from flow-sorted naive CD8^+^ T cells (n = 300 per condition) using a NucleoSpin RNA XS Kit (Macherey-Nagel), and cDNA was synthesized using Reverse Transcription Master Mix (Fluidigm). Specific targets were amplified using PreAmp Master Mix (Fluidigm). Gene expression was assessed using a BioMark HD System (Fluidigm) with EvaGreen Supermix (Bio-Rad). RNA expression levels were calculated using the 2^−ΔΔCT^ method with reference to a housekeeping gene (human 18S) [16].

### Statistics

Univariate statistical analyses were performed using nonparametric tests in Prism software version 8 (GraphPad Software Inc.). Significance was assigned at p < 0.05.

### Study Approval

Ethical approval was granted by the Comité de Protection des Personnes of the Pitié Salpétrière Hospital (Paris, France). All volunteers provided written informed consent in accordance with the Declaration of Helsinki.

## RESULTS

### Naive CD8^+^ T cells in the elderly exhibit altered basal activation status and proliferative capacity

We previously demonstrated that naïve CD8^+^ T cells from elderly individuals generate, upon encountering their cognate antigen, qualitatively altered effector cells characterized by a skewed differentiation status and a poor cytotoxic potential [5]. To better characterize this phenomenon, we compared the activation profiles of naive CD8^+^ T cells in middle-aged and elderly individuals, mimicking antigen-driven signals with plate-bound anti-CD3. No age-related differences in activation *per se* were detected 24 hr after stimulation, as determined by measuring the upregulation of CD69 and CD134 (Figure 1a). Despite these phenotypic similarities, naive CD8^+^ T cells from elderly individuals proliferated to a lower extent than naive CD8^+^ T cells from middle-aged individuals in response to stimulation with plate-bound anti-CD3 (Figure 1b), consistent with their poor expansion upon priming [5]. Susceptibility to apoptosis upon activation was also more commonly observed among activated naive CD8^+^ T cells from elderly *versus* middle-aged individuals, as determined by the intracellular expression of active caspase-3, known to be a central element of the apoptotic pathway (Figure 1c). Of note, the proliferative capacity and susceptibility to apoptosis of naive CD8^+^ T cells upon stimulation were inversely correlated, irrespective of age (Figure 1d).

**Figure 1.**
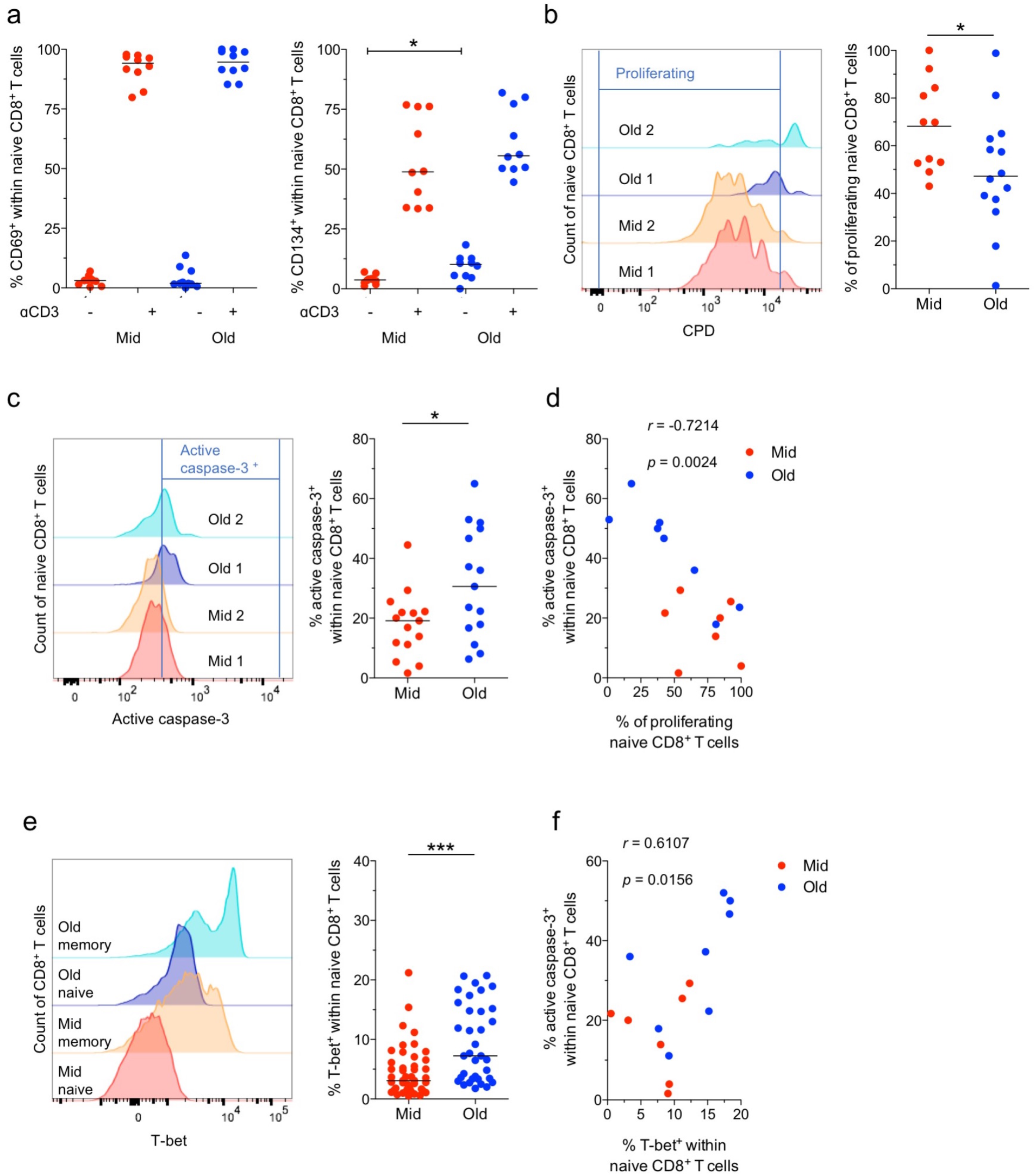
Activation and proliferation of naive CD8^+^ T cells. (**a - c**) PBMCs from middle-aged (Mid - 19 to 55 years old) and elderly individuals (Old - 65 to 95 years old) were incubated in the absence or presence of plate-bound anti-CD3. Surface expression of the activation markers CD69 and CD134 was measured after 24 hr (a), proliferation was measured after 4 days (b), and active caspase-3 expression was measured after 1 day (c). Left panels: representative flow cytometry profiles. Right panels: data summaries. Data are shown for naive CD8^+^ T cells. Each dot represents one donor. Horizontal lines indicate median values. * p < 0.05 (Mann-Whitney U test). (**d**) Correlation between the frequency of naive CD8^+^ T cells that proliferated after stimulation and the frequency of naive CD8^+^ T cells that expressed active caspase-3 after stimulation. Each dot represents one donor. Significance was determined using Spearman’s rank correlation. (**e**) T-bet expression was measured in unstimulated naive CD8^+^ T cells from middle-aged (Mid) and elderly individuals (Old). Left panel: representative flow cytometry profiles. Right panel: data summary. Each dot represents one donor. Horizontal lines indicate median values. *** p < 0.001 (Mann-Whitney *U* test). (**f**) Correlation between the basal expression frequency of T-bet and the activation-induced expression frequency of active caspase-3 among naive CD8^+^ T cells. Each dot represents one donor. Significance was determined using Spearman’s rank correlation.

Unstimulated naive CD8^+^ T cells from elderly individuals expressed CD134 more commonly than unstimulated naive CD8^+^ T cells from middle-aged individuals, suggesting increased levels of basal activation with advanced age (Figure 1a). To consolidate this observation, we measured the expression of T-bet, which is classically upregulated in response to activation. The basal expression frequencies of T-bet mirrored the basal expression frequencies of CD134 (Figure 1e). Equivalent results were obtained using a more stringent definition of naive CD8^+^ T cells (Figures S1a and S1b), which excluded phenotypically similar memory CD8^+^ T cells [17]. The basal expression frequency of T-bet also correlated directly with the activation-induced expression frequency of active caspase-3 among naive CD8^+^ T cells (Figure 1f).

Collectively, these data indicate that the deficit in proliferation of old naive CD8^+^ T cells is associated with higher basal activation status and susceptibility to apoptosis, despite a largely unaltered early response to activation.

### Naive CD8^+^ T cells in the elderly are metabolically distinct

Signals transduced via the TCR elicit an mTOR-driven metabolic switch that supports the function and the viability of activated naive CD8^+^ T cells [9]. We therefore assessed mTOR activity in parallel by quantifying phospho-S6 (pS6). In line with the comparable activation profiles, naive CD8^+^ T cells in middle-aged and elderly individuals upregulated mTOR activity to a similar extent after stimulation with plate-bound anti-CD3 (Figure 2a). To extend these findings, we measured the expression of metabolism-related genes, comparing unstimulated and activated naive CD8^+^ T cells (Figure 2b). Genes encoding various enzymes involved in glycolysis were upregulated similarly in flow-sorted naive CD8^+^ T cells from middle-aged and elderly individuals after stimulation with plate-bound anti-CD3. In contrast, genes associated with lipid metabolism or signaling pathways involved in metabolic regulation were not generally upregulated in response to stimulation, with the exception of *MYC*, which was overexpressed in activated naive CD8^+^ T cells, irrespective of age. Genes that play a critical role in the metabolic switch were also overexpressed in activated naive CD8^+^ T cells, irrespective of age, with the exception of *HIF1* and *RPS6KB1*, which were upregulated to a greater extent in activated naive CD8^+^ T cells from middle-aged *versus* elderly individuals. This dataset indicates a rather intact although suboptimal metabolic switch, with a physiological mTOR activation, in naive CD8^+^ T cells of elderly subjects upon TCR ligation.

**Figure 2.**
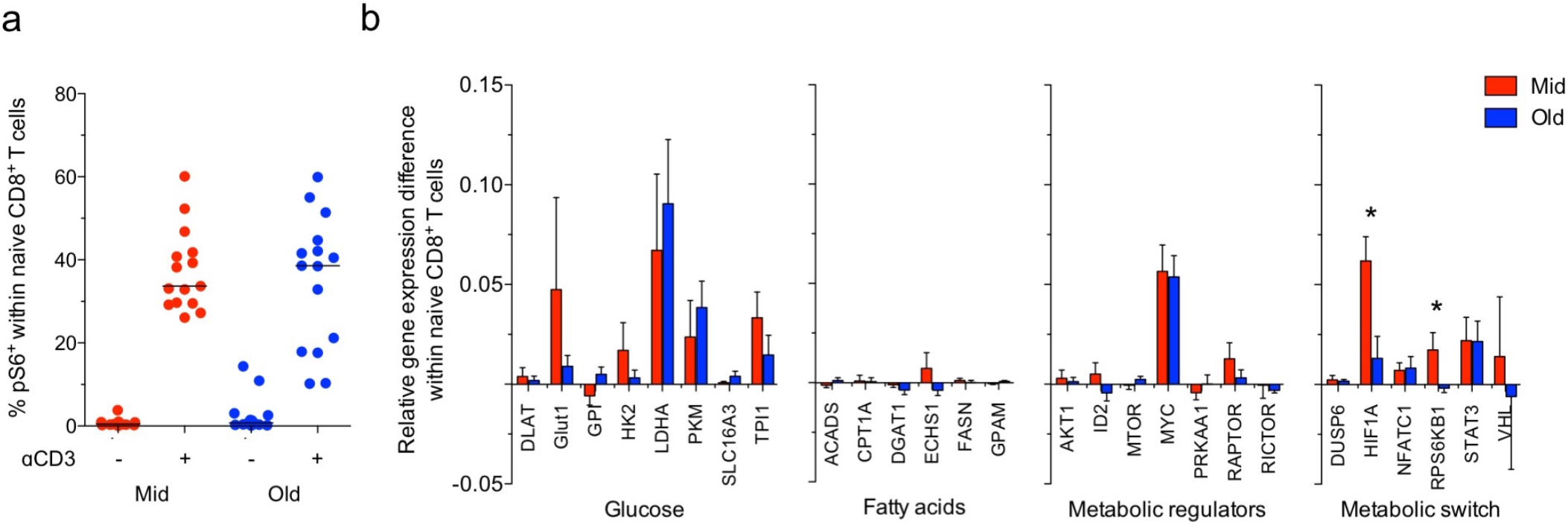
Metabolic switch in the naive CD8^+^ T cell compartment. (**a**) PBMCs from middle-aged (Mid) and elderly individuals (Old) were incubated in the absence or presence of plate-bound anti-CD3. Intracellular expression of mTOR activity marker pS6 was measured after 3 hr. Data are shown for naive CD8^+^ T cells. Each dot represents one donor. Horizontal lines indicate median values. * p < 0.05 (Mann-Whitney *U* test). (**b**) Flow-sorted naive CD8^+^ T cells from middle-aged (n = 4; Mid) and elderly individuals (n = 5; Old) were incubated in the absence or presence of plate-bound anti-CD3. Gene expression levels were measured after 5 hr. Data are shown relative to the unstimulated condition. Bars indicate mean ± SEM. * p < 0.05 (Mann-Whitney *U* test).

To explore the age-related homeostatic change in more detail, i.e. prior to activation, we investigated the biochemical features of quiescent naive CD8^+^ T cells. These cells are present at very low frequencies in elderly donors [5], posing challenges to the determination of metabolic fluxes, for instance using standard approaches like Seahorse technology. We therefore engaged in the characterization of different metabolic pathways using flow cytometry and gene expression profiles. Glycolysis is the main metabolic pathway that supports the activation of naive CD8^+^ T cells [9,18]. We found no significant differences in basal glucose uptake between unstimulated naive CD8^+^ T cells from middle-aged individuals and unstimulated naive CD8^+^ T cells from elderly individuals (Figure 3a). Moreover, we found similar basal expression levels of glycolysis-related genes, with the exception of *HK2*, which was overexpressed in unstimulated naive CD8^+^ T cells from middle-aged *versus* elderly individuals (Figure 3b). This gene encodes a selectively regulated isoform of hexokinase [19], which catalyzes glucose phosphorylation, and is usually induced in response to stimulation via the TCR [19,20]. Interestingly, fatty acid (FA) uptake was decreased among unstimulated naive CD8^+^ T cells from middle-aged *versus* elderly individuals (Figure 3c), but this difference was not associated with significant changes in the expression levels of genes encoding various enzymes involved in FA oxidation (FAO) or FA synthesis (FAS). However, we noted that *DGAT1*, which encodes diacylglycerol O-acyltransferase 1, a key enzyme involved in the storage of FAs as triacylglycerol (TAG), was expressed at lower levels, although not statistically significant, in unstimulated naive CD8^+^ T cells from middle-aged *versus* elderly individuals (Figure 3d).

**Figure 3.**
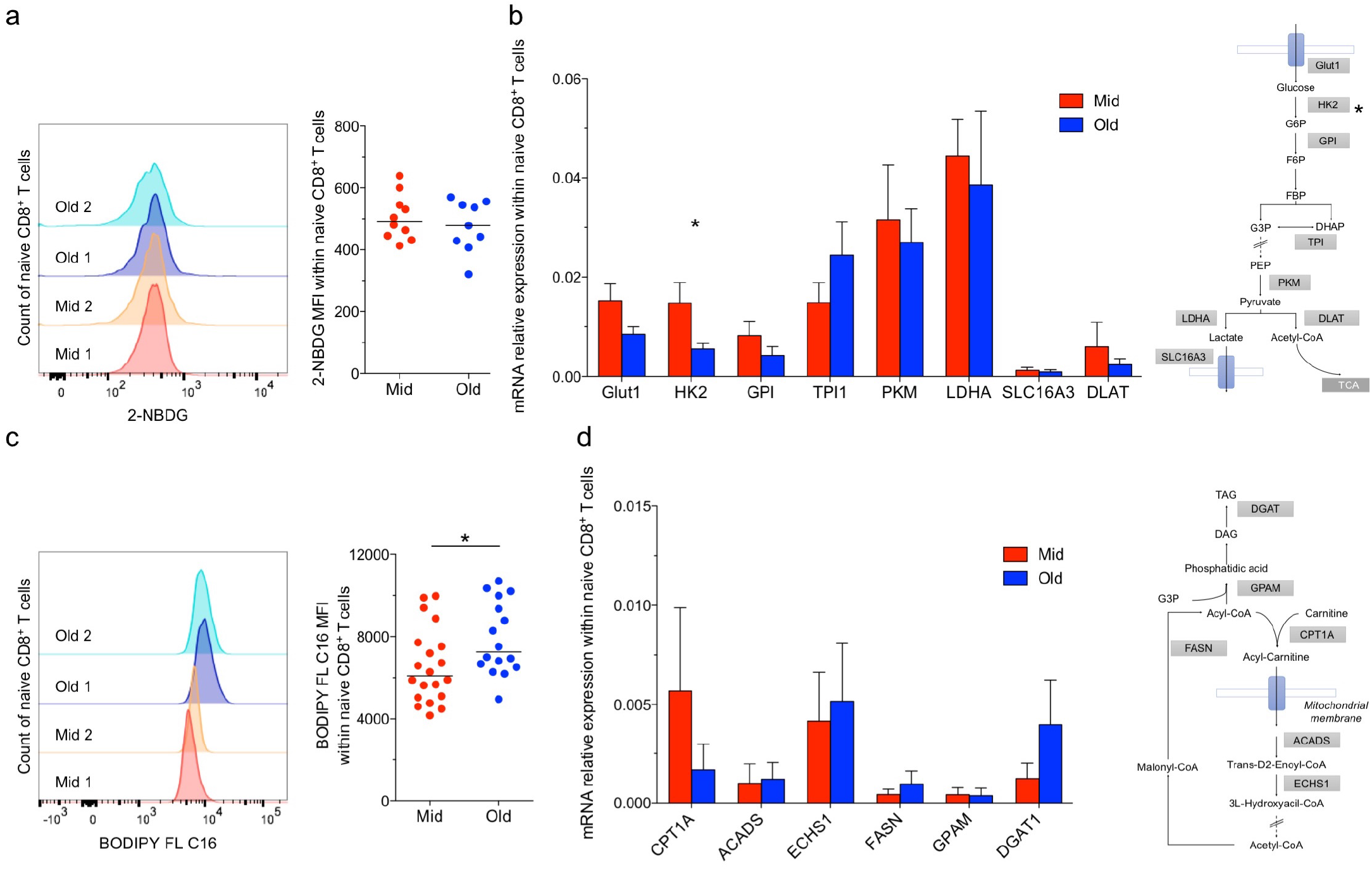
Basal metabolism in the naive CD8^+^ T cell compartment. (**a** & **c**) Glucose (a) and FA uptake (c) were measured in unstimulated naive CD8^+^ T cells from middle-aged (Mid) and elderly individuals (Old) by determining the mean fluorescence intensity (MFI) of 2-NBDG, and BODIPY FL C16, respectively. Left panels: representative flow cytometry profiles. Right panels: data summaries. Each dot represents one donor. Horizontal lines indicate median values. * p < 0.05 (Mann-Whitney *U* test). (**b** & **d**) Expression levels of genes related to glucose (b) and fatty acid metabolism (d) were measured in unstimulated naive CD8^+^ T cells flow-sorted from middle-aged (n = 5; Mid) and elderly individuals (n = 5; Old). Bars indicate mean ± SEM. * p < 0.05 (Mann-Whitney *U* test).

In addition, we observed that unstimulated naive CD8^+^ T cells from middle-aged individuals stored lower amounts of neutral lipids (NLs) than unstimulated naive CD8^+^ T cells from elderly individuals (Figure 4a). In further experiments, we assessed the basal expression levels of various genes encoding transcription factors involved in metabolic regulation. Similar patterns of expression were observed in unstimulated naive CD8^+^ T cells, irrespective of age, with the exception of *ID2*, which was expressed at lower levels in unstimulated naive CD8^+^ T cells from middle-aged *versus* elderly individuals (Figure 4b). *ID2* is involved in metabolic adaptation [21,22] and promotes lipid storage via the downmodulation of *PGC-1α* [22], which enhances FAO and inhibits TAG synthesis [23]. Moreover, *ID2* promotes an overall increase in mitochondrial membrane potential (ΔΨM), without affecting mitochondrial biogenesis or, by extension, mitochondrial mass [21]. In line with these known functions, ΔΨM was lower in unstimulated naive CD8^+^ T cells from middle-aged *versus* elderly individuals (Figure 4c), whereas mitochondrial mass was largely unaffected by age (Figure 4d). We also noted a direct correlation between ΔΨM and the frequency of unstimulated naive CD8^+^ T cells that expressed T-bet, suggesting a link with the loss of quiescence (Figure S2).

**Figure 4.**
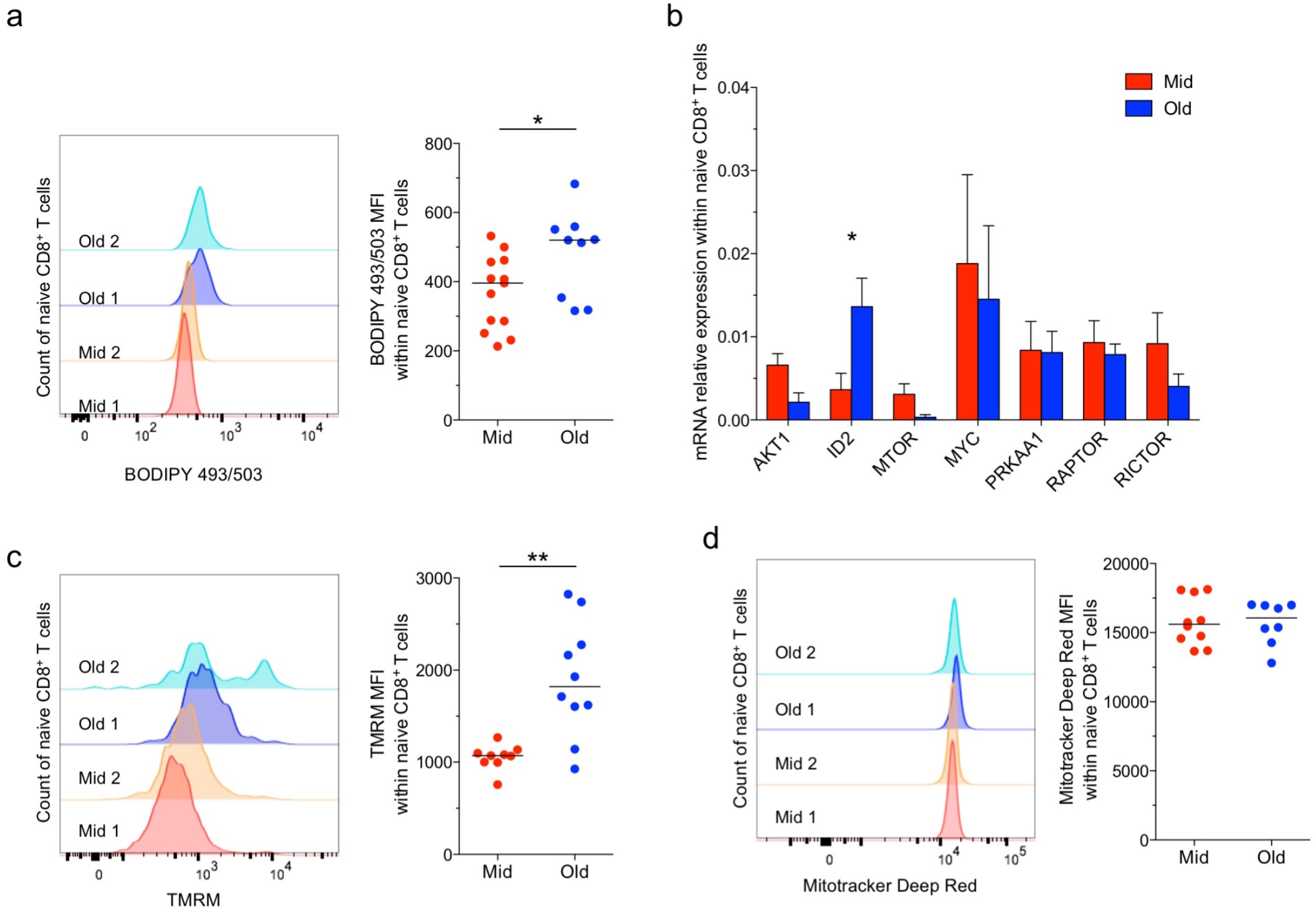
Metabolic control in the naive CD8^+^ T cell compartment. (**a, c** & **d**) NL content (a) and mitochondrial membrane potential (C) and mass (D) were measured in unstimulated naive CD8^+^ T cells from middle-aged (Mid) and elderly individuals (Old) by determining the mean fluorescence intensity (MFI) of BODIPY 493/503, TMRM and Mitotracker Deep Red, respectively. Left panels: representative flow cytometry profiles. Right panels: data summaries. Each dot represents one donor. Horizontal lines indicate median values. * p < 0.05, ** p < 0.01 (Mann-Whitney *U* test). (**b**) Expression levels of genes related to signaling pathways involved in metabolic regulation were measured in unstimulated naive CD8^+^ T cells flow-sorted from middle-aged (n = 5; Mid) and elderly individuals (n = 5; Old). Bars indicate mean ± SEM. * p < 0.05 (Mann-Whitney *U* test).

Collectively, these data revealed an age-related shift in the basal metabolic properties of naive CD8^+^ T cells, typified by a supranormal ΔΨM and high levels of FA uptake and NL storage.

### Naive CD8^+^ T cells in the elderly can be reinvigorated with lipid-altering drugs

While FAs are indispensable for cellular lifespan, and in particular for the formation of cell membranes, their excessive levels can have negative effects on cell physiology. T cell homeostasis and viability can for instance be affected by high levels of FAs [24,25].We therefore assessed the potential link between the functional properties of old naïve T cells and their altered lipid metabolism. We first observed direct correlations between the frequency of unstimulated naive CD8^+^ T cells that expressed active caspase-3 and basal levels of FA uptake (Figure 5a) and NL content (Figure 5b). To determine the biological relevance of these associations, we treated naive CD8^+^ T cells with rosiglitazone, a drug known to foster lipid catabolism by activating triglyceride lipase [26] and preventing the conversion of FAs into NLs [27]. Exposure to rosiglitazone in serum starvation conditions boosted indeed NL catabolism in naive CD8^+^ T cells, as observed with a decrease of cellular NL content (Figure 5c). Pretreatment with rosiglitazone reduced activation-induced active caspase-3 expression among naive CD8^+^ T cells of elderly subjects (Figure 5d). Similar results were obtained using fenofibrate, which induces lipid catabolism by enhancing FAO (Figure S3).

**Figure 5.**
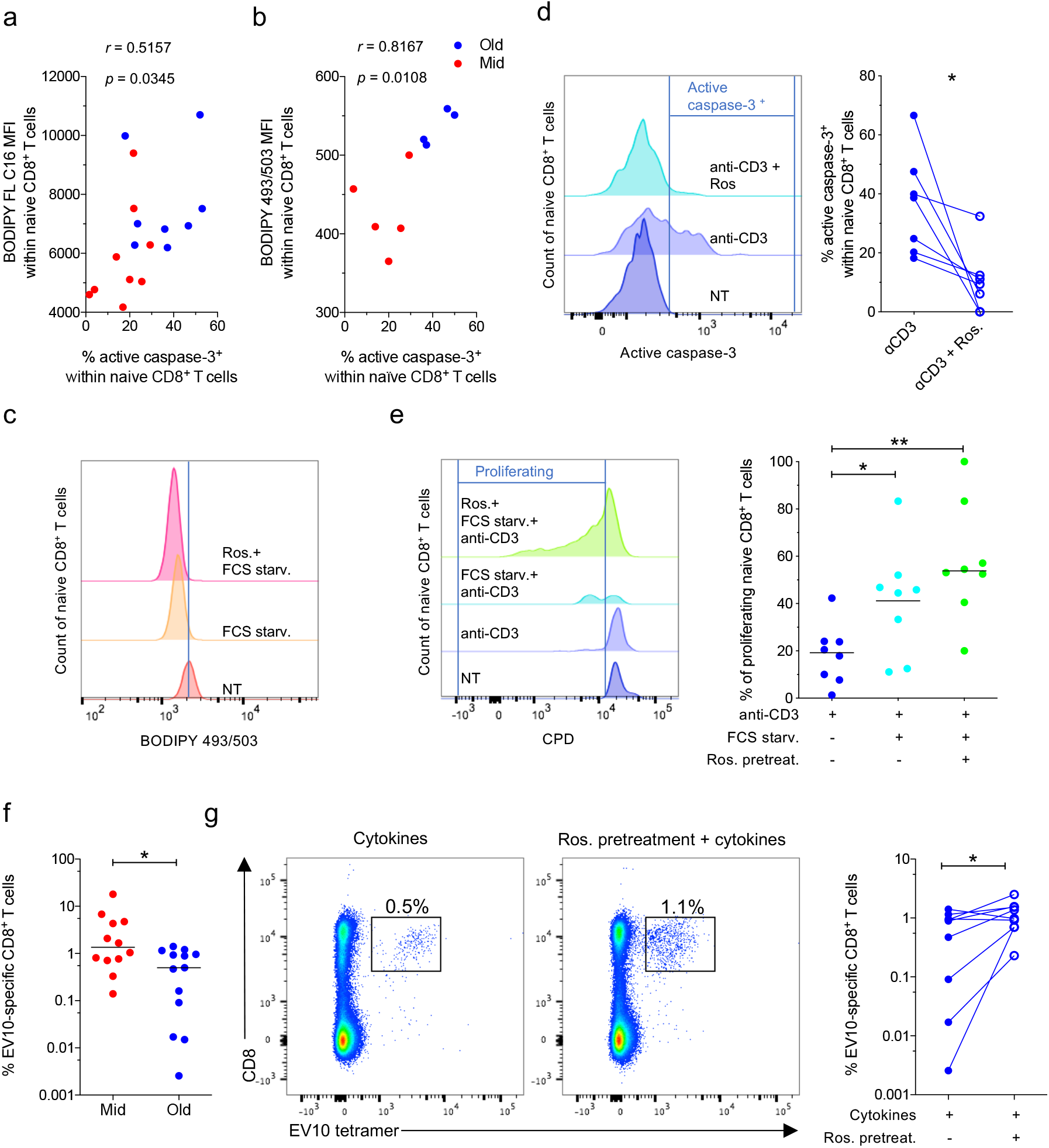
Effects of lipid-altering drugs in the naive CD8^+^ T cell compartment. (**a** & **b**) Correlations between the frequency of unstimulated naive CD8^+^ T cells that expressed active caspase-3 and basal levels of FA uptake (a) and NL content (b) measured by determining the mean fluorescence intensity (MFI) of BODIPY FL C16 and BODIPY 493/503, respectively. Each dot represents one donor. Significance was determined using Spearman’s rank correlation. (**c**) PBMCs were preincubated for 2 days in serum-free medium in the absence or presence of rosiglitazone (Ros). NL content was measured in naive CD8^+^ T cells by determining the mean fluorescence intensity (MFI) of BODIPY 493/503. Flow cytometry profiles are representative of five independent experiments. (**d**) PBMCs from elderly individuals (n = 8) were stimulated with plate-bound anti-CD3 in the absence or presence of rosiglitazone (Ros). Active caspase-3 expression was measured after 24 hr. Data are shown for naive CD8^+^ T cells. Left panel: representative flow cytometry profiles. Right panel: data summary. Bars indicate mean ± SEM. * p < 0.05 (Wilcoxon signed rank test). (**e**) PBMCs from elderly individuals were preincubated for 2 days in serum-free medium in the absence or presence of rosiglitazone (Ros) and stimulated with plate-bound anti-CD3. Proliferation was measured after 4 days. Data are shown for naive CD8^+^ T cells. Left panel: representative flow cytometry profiles. Right panel: data summary. Each dot represents one donor. Horizontal lines indicate median values. * p < 0.05, ** p < 0.01 (Mann-Whitney *U* test). (**f**) Percentage of tetramer^+^ EV10-specific CD8^+^ T cells expanded for 10 days in the presence of Flt3 ligand and a cocktail of inflammatory cytokines. Each dot represents one donor. Horizontal lines indicate median values. * p < 0.05 (Mann-Whitney *U* test). (**g**) Percentage of tetramer^+^ EV10-specific CD8^+^ T cells of elderly individuals expanded for 10 days in the presence of Flt3 ligand and a cocktail of inflammatory cytokines after preincubation for 2 days in the absence or presence of rosiglitazone (Ros). Left panel: representative flow cytometry profiles. Right panel: data summary. Each dot represents one donor. Horizontal lines indicate median values. * p < 0.05 (Wilcoxon signed rank test). NT: not treated.

With regards to proliferative capacity, naive CD8^+^ T cells from elderly individuals proliferated to a greater extent after serum starvation, and the addition of rosiglitazone further enhanced these activation-induced proliferative responses (Figure 5e). To assess the potential relevance of these findings in the context of antigen-driven immune responses, we used a previously established *in vitro* model to prime naive CD8^+^ T cells specific for EV10, a melanoma-derived HLA-A2 restricted epitope, starting from PBMCs of melanoma-naïve individuals. We developed and have used this assay to study human CD8^+^ T cell priming [5]. As expected from the proliferation data, EV10-specific CD8^+^ T cells from middle-aged individuals expanded to a greater extent than EV10-specific CD8^+^ T cells from elderly individuals (Figure 5f), and preincubation with rosiglitazone consistently enhanced the expansion of EV10-specific CD8^+^ T cells from elderly individuals (Figure 5g).

Collectively, these data indicate that age-related functional deficits associated with abnormal lipid metabolism and greater levels of basal activation in the naive CD8^+^ T cell compartment may be, at least in part, reversed in the presence of rosiglitazone, opening potentially useful therapeutic approaches to enhance immune reactivity against newly encountered antigens in the elderly.

## DISCUSSION

A detailed understanding of age-related deficits in the naive T cell compartment is essential for the rational development of immunotherapies and vaccines that protect elderly individuals from emerging threats. We found that naive CD8^+^ T cells in elderly individuals were susceptible to apoptosis and proliferated suboptimally in response to stimulation. These abnormalities were associated with enhanced levels of basal activation, measured in terms of ΔΨM and the *ex vivo* expression frequencies of T-bet and CD134.

Recent studies have shown that metabolic processes govern the behavior of T cells [9,18,29,30]. In the naive CD8^+^ T cell compartment, autophagy and glycolysis are typically upregulated in response to activation [9,31,32], whereas homeostatic energy requirements are fulfilled primarily via FAO [11,33–36]. We observed that, at rest, naive CD8^+^ T cells in the elderly displayed abnormally high levels of FA uptake and stored abnormally high amounts of NLs. In line with previous reports showing that high levels of intracellular lipids may be toxic [9,24,37], we found that active caspase-3 expression correlated directly with FA uptake and NL content in the naive CD8^+^ T cell compartment. Lipids are essential for T cell activation and proliferation [28]. Excessively high amounts of intracellular lipids can nonetheless impair T cell proliferation and viability [38–40]. Accordingly, we found that activation-induced initiation of the apoptotic pathway was reduced by interventions enhancing lipid clearance in naive CD8^+^ T cells. Of note, NLs *per se* are not toxic. The conversion of FAs into NLs therefore most likely protects against lipotoxicity under homeostatic conditions [24].

The heightened basal activation status of naive CD8^+^ T cells in elderly individuals seemed to be sustained energetically by increased mitochondrial activity, given that T-bet expression correlated directly with ΔΨM. Inflammation is closely linked with metabolic dysregulation in the elderly [41]. High levels of circulating proinflammatory cytokines and lipids are common features of advanced age and may contribute to the disruption of cellular quiescence. Moreover, hematopoietic progenitor cells in elderly individuals are often metabolically active, and this characteristic may be heritable [42]. Increased rates of homeostatic proliferation are required to maintain naive CD8^+^ T cell numbers in the elderly [43], and the predominant energetic pathway that supports this process is thought to be FAO [44,45]. It is therefore plausible that high basal levels of FA uptake and NL storage constitute a bioenergetic pattern that favors homeostatic proliferation.

Aging is characterized by profound metabolic perturbations [46], including increased lipogenesis [47] and reduced lipolysis [48], leading to higher systemic levels of free FAs and TAG [41,49]. The combination of a homeostatic environment and high systemic levels of proinflammatory cytokines and lipids may potentially underlie the altered metabolism and functional deficits that characterize naive CD8^+^ T cells in the elderly. A key finding of our study was the observation that rosiglitazone, known to foster lipid catabolism, can improve these abnormalities, and enhance old naïve T cell priming. The experimental model we exploited (i.e. *in vitro* priming of melanoma-specific naïve CD8^+^ T cells) was previously shown to mimic *in vivo* responses in both mice and humans [5,50,51]. The positive effect of rosiglitazone in this model is an encouraging result from a translational perspective. Of note, a recent study showed that treatment with this drug was associated with a delay of age□associated metabolic disease and extend longevity in old mice [52]. Although rosiglitazone, initially licensed for the treatment of diabetes mellitus, was eventually withdrawn due several side effects, it serves as proof of concept that inducing lipolysis prior to T-cell priming could be beneficial in the context of vaccinations and immunotherapy, akin to recent data showing that increased FAO can improve the development and the quality of effector CD8^+^ T cells [9,53–55]. Further studies are therefore warranted to screen whether other lipid-altering drugs may be useful adjunctive therapies to enhance adaptive immune responses against previously unencountered antigens, particularly in elderly individuals, who often respond poorly to vaccination and remain vulnerable to emerging pathogens, such as seasonal influenza viruses and SARS-CoV-2.

## Supporting information

Supp Mat

## Contributors

F.N., R.G., and V.A. conceptualized the project; F.N., M.P.C.P., L.P., E.Ga., V.F., and J.J.F. performed experiments and analyzed data; M.D., E.C., H.V., E.Go., S.L.L., D.A.P., A.T., and J.B. provided critical resources; F.N., J.F.F., R.G., and V.A. drafted the manuscript; F.N., D.A.P., A.C., R.G., and V.A. edited the manuscript; D.A.P., A.T., A.C., and V.A. acquired funds to support the work. All authors contributed intellectually and approved the manuscript.

## Declaration of Competing Interests

The authors declare no conflict of interest.

## Acknowledgements

We thank Simone Candioli and Silvio Spatocco (University of Ferrara, Italy), Irene Bonazzi and Ilaria Signoretto (University of Padua, Italy), and Alain Savenay (AP-HP, Paris, France) for technical assistance, Mario Pende (Institut Necker-Enfants Malades, Paris, France) for helpful discussions, and Veronique Morin and Rima Zoorob (INSERM U1135, Paris, France) and Silvia Menegatti and Lars Rogge (Institut Pasteur, Paris, France) for assistance with gene expression analyses. This work was supported by the Agence National de la Recherche (ANR-14-CE14-0030-01) and the Université Franco-Italienne/Università Italo-Francese (Galileo Project G10-718; PHC Galilee Project 39582TJ). Core provision of the FACSCanto II was funded by the University of Ferrara (Bando per l’acquisizione di strumenti per la ricerca di ateneo-2015). D.A.P. was supported by a Wellcome Trust Senior Investigator Award (100326/Z/12/Z).

## Data Sharing

Data collected for the study will be made available upon reasonable request made to the corresponding authors FN and VA.

## Notes

### Competing Interest Statement

The authors have declared no competing interest.

## REFERENCES

[1] Nicoli F, Solis-Soto MT, Paudel D, et al. Age-related decline of de novo T cell responsiveness as a cause of COVID-19 severity. Geroscience. 2020; 42: 1015–9.

[2] Dorshkind K, Swain S. Age-associated declines in immune system development and function: causes, consequences, and reversal. Curr Opin Immunol. 2009; 21: 404–7.

[3] Ventura MT, Casciaro M, Gangemi S, Buquicchio R. Immunosenescence in aging: between immune cells depletion and cytokines up-regulation. Clin Mol Allergy. 2017; 15: 21.

[4] Fulop T, Larbi A, Dupuis G, et al. Immunosenescence and Inflamm-Aging As Two Sides of the Same Coin: Friends or Foes? Front Immunol. 2017; 8: 1960.

[5] Briceno O, Lissina A, Wanke K, et al. Reduced naive CD8(+) T-cell priming efficacy in elderly adults. Aging Cell. 2016; 15: 14–21.

[6] Nikolich-Zugich J. Aging of the T cell compartment in mice and humans: from no naive expectations to foggy memories. J Immunol. 2014; 193: 2622–9.

[7] Wertheimer AM, Bennett MS, Park B, et al. Aging and cytomegalovirus infection differentially and jointly affect distinct circulating T cell subsets in humans. J Immunol. 2014; 192: 2143–55.

[8] Palmer CS, Ostrowski M, Balderson B, Christian N, Crowe SM. Glucose metabolism regulates T cell activation, differentiation, and functions. Front Immunol. 2015; 6: 1.

[9] Nicoli F, Papagno L, Frere JJ, et al. Naive CD8(+) T-Cells Engage a Versatile Metabolic Program Upon Activation in Humans and Differ Energetically From Memory CD8(+) T-Cells. Front Immunol. 2018; 9: 2736.

[10] Nicoli F. Angry, Hungry T-Cells: How Are T-Cell Responses Induced in Low Nutrient Conditions? Immunometabolism. 2020; 2: e200004.

[11] Almeida L, Lochner M, Berod L, Sparwasser T. Metabolic pathways in T cell activation and lineage differentiation. Semin Immunol. 2016; 28: 514–24.

[12] Baylis D, Bartlett DB, Patel HP, Roberts HC. Understanding how we age: insights into inflammaging. Longev Healthspan. 2013; 2: 8.

[13] Price DA, Brenchley JM, Ruff LE, et al. Avidity for antigen shapes clonal dominance in CD8+ T cell populations specific for persistent DNA viruses. J Exp Med. 2005; 202: 1349–61.

[14] Lissina A, Briceno O, Afonso G, et al. Priming of Qualitatively Superior Human Effector CD8+ T Cells Using TLR8 Ligand Combined with FLT3 Ligand. J Immunol. 2016; 196: 256–63.

[15] Alanio C, Nicoli F, Sultanik P, et al. Bystander hyperactivation of preimmune CD8+ T cells in chronic HCV patients. Elife. 2015; 4: 1–20.

[16] Nicoli F, Gallerani E, Sforza F, et al. The HIV-1 Tat protein affects human CD4+ T-cell programing and activation, and favors the differentiation of naive CD4+ T cells. AIDS. 2018; 32: 575–81.

[17] Pulko V, Davies JS, Martinez C, et al. Human memory T cells with a naive phenotype accumulate with aging and respond to persistent viruses. Nat Immunol. 2016; 17: 966–75.

[18] Zhang L, Romero P. Metabolic Control of CD8(+) T Cell Fate Decisions and Antitumor Immunity. Trends Mol Med. 2018; 24: 30–48.

[19] Tan H, Yang K, Li Y, et al. Integrative Proteomics and Phosphoproteomics Profiling Reveals Dynamic Signaling Networks and Bioenergetics Pathways Underlying T Cell Activation. Immunity. 2017; 46: 488–503.

[20] Lis P, Dylag M, Niedzwiecka K, et al. The HK2 Dependent “Warburg Effect” and Mitochondrial Oxidative Phosphorylation in Cancer: Targets for Effective Therapy with 3-Bromopyruvate. Molecules. 2016; 21.

[21] Zhang Z, Rahme GJ, Chatterjee PD, Havrda MC, Israel MA. ID2 promotes survival of glioblastoma cells during metabolic stress by regulating mitochondrial function. Cell Death Dis. 2017; 8: e2615.

[22] Hou TY, Ward SM, Murad JM, Watson NP, Israel MA, Duffield GE. ID2 (inhibitor of DNA binding 2) is a rhythmically expressed transcriptional repressor required for circadian clock output in mouse liver. J Biol Chem. 2009; 284: 31735–45.

[23] Morris EM, Meers GM, Booth FW, et al. PGC-1alpha overexpression results in increased hepatic fatty acid oxidation with reduced triacylglycerol accumulation and secretion. Am J Physiol Gastrointest Liver Physiol. 2012; 303: G979–92.

[24] de Jong AJ, Kloppenburg M, Toes RE, Ioan-Facsinay A. Fatty acids, lipid mediators, and T-cell function. Front Immunol. 2014; 5: 483.

[25] Nicoli F, Paul S, Appay V. Harnessing the Induction of CD8+ T-Cell Responses Through Metabolic Regulation by Pathogen-Recognition-Receptor Triggering in Antigen Presenting Cells. Frontiers in Immunology. 2018; 9.

[26] Kershaw EE, Schupp M, Guan HP, Gardner NP, Lazar MA, Flier JS. PPARgamma regulates adipose triglyceride lipase in adipocytes in vitro and in vivo. Am J Physiol Endocrinol Metab. 2007; 293: E1736–45.

[27] Askari B, Kanter JE, Sherrid AM, et al. Rosiglitazone inhibits acyl-CoA synthetase activity and fatty acid partitioning to diacylglycerol and triacylglycerol via a peroxisome proliferator-activated receptor-gamma-independent mechanism in human arterial smooth muscle cells and macrophages. Diabetes. 2007; 56: 1143–52.

[28] Angela M, Endo Y, Asou HK, et al. Fatty acid metabolic reprogramming via mTOR-mediated inductions of PPARgamma directs early activation of T cells. Nat Commun. 2016; 7: 13683.

[29] Gubser PM, Bantug GR, Razik L, et al. Rapid effector function of memory CD8+ T cells requires an immediate-early glycolytic switch. Nat Immunol. 2013; 14: 1064–72.

[30] O’Neill LA, Kishton RJ, Rathmell J. A guide to immunometabolism for immunologists. Nat Rev Immunol. 2016; 16: 553–65.

[31] Arnold CR, Pritz T, Brunner S, et al. T cell receptor-mediated activation is a potent inducer of macroautophagy in human CD8(+)CD28(+) T cells but not in CD8(+)CD28(-) T cells. Exp Gerontol. 2014; 54: 75–83.

[32] Whang MI, Tavares RM, Benjamin DI, et al. The Ubiquitin Binding Protein TAX1BP1 Mediates Autophagasome Induction and the Metabolic Transition of Activated T Cells. Immunity. 2017; 46: 405–20.

[33] O’Sullivan D, van der Windt GJ, Huang SC, et al. Memory CD8(+) T cells use cell-intrinsic lipolysis to support the metabolic programming necessary for development. Immunity. 2014; 41: 75–88.

[34] Green WD, Beck MA. Obesity altered T cell metabolism and the response to infection. Curr Opin Immunol. 2017; 46: 1–7.

[35] Pearce EL, Walsh MC, Cejas PJ, et al. Enhancing CD8 T-cell memory by modulating fatty acid metabolism. Nature. 2009; 460: 103–7.

[36] Raud B, McGuire PJ, Jones RG, Sparwasser T, Berod L. Fatty acid metabolism in CD8(+) T cell memory: Challenging current concepts. Immunol Rev. 2018; 283: 213–31.

[37] Zurier RB, Rossetti RG, Seiler CM, Laposata M. Human peripheral blood T lymphocyte proliferation after activation of the T cell receptor: effects of unsaturated fatty acids. Prostaglandins Leukot Essent Fatty Acids. 1999; 60: 371–5.

[38] Takahashi HK, Cambiaghi TD, Luchessi AD, et al. Activation of survival and apoptotic signaling pathways in lymphocytes exposed to palmitic acid. J Cell Physiol. 2012; 227: 339–50.

[39] Fernanda Cury-Boaventura M, Cristine Kanunfre C, Gorjao R, Martins de Lima T, Curi R. Mechanisms involved in Jurkat cell death induced by oleic and linoleic acids. Clin Nutr. 2006; 25: 1004–14.

[40] Howie D, Cobbold SP, Adams E, et al. Foxp3 drives oxidative phosphorylation and protection from lipotoxicity. JCI Insight. 2017; 2: e89160.

[41] Pararasa C, Ikwuobe J, Shigdar S, et al. Age-associated changes in long-chain fatty acid profile during healthy aging promote pro-inflammatory monocyte polarization via PPARgamma. Aging Cell. 2016; 15: 128–39.

[42] Fali T, Fabre-Mersseman V, Yamamoto T, et al. Elderly human hematopoietic progenitor cells express cellular senescence markers and are more susceptible to pyroptosis. JCI Insight. 2018; 3.

[43] Sauce D, Larsen M, Fastenackels S, et al. Lymphopenia-driven homeostatic regulation of naive T cells in elderly and thymectomized young adults. J Immunol. 2012; 189: 5541–8.

[44] Chang CH, Curtis JD, Maggi LB, Jr., et al. Posttranscriptional control of T cell effector function by aerobic glycolysis. Cell. 2013; 153: 1239–51.

[45] Ibitokou SA, Dillon BE, Sinha M, et al. Early Inhibition of Fatty Acid Synthesis Reduces Generation of Memory Precursor Effector T Cells in Chronic Infection. J Immunol. 2018; 200: 643–56.

[46] Bonomini F, Rodella LF, Rezzani R. Metabolic syndrome, aging and involvement of oxidative stress. Aging Dis. 2015; 6: 109–20.

[47] Kuhla A, Blei T, Jaster R, Vollmar B. Aging is associated with a shift of fatty metabolism toward lipogenesis. J Gerontol A Biol Sci Med Sci. 2011; 66: 1192–200.

[48] Toth MJ, Tchernof A. Lipid metabolism in the elderly. Eur J Clin Nutr. 2000; 54 Suppl 3: S121–5.

[49] Mc Auley MT, Mooney KM. Computationally Modeling Lipid Metabolism and Aging: A Mini-review. Comput Struct Biotechnol J. 2015; 13: 38–46.

[50] Gutjahr A, Papagno L, Nicoli F, et al. The STING ligand cGAMP potentiates the efficacy of vaccine-induced CD8+ T cells. JCI Insight. 2019; 4.

[51] Gutjahr A, Papagno L, Nicoli F, et al. Cutting Edge: A Dual TLR2 and TLR7 Ligand Induces Highly Potent Humoral and Cell-Mediated Immune Responses. J Immunol. 2017; 198: 4205–9.

[52] Xu L, Ma X, Verma N, et al. PPARgamma agonists delay age-associated metabolic disease and extend longevity. Aging Cell. 2020; 19: e13267.

[53] Zhang Y, Kurupati R, Liu L, et al. Enhancing CD8+ T Cell Fatty Acid Catabolism within a Metabolically Challenging Tumor Microenvironment Increases the Efficacy of Melanoma Immunotherapy. Cancer Cell. 2017; 32: 377–91 e9.

[54] Chowdhury PS, Chamoto K, Honjo T. Combination therapy strategies for improving PD-1 blockade efficacy: a new era in cancer immunotherapy. J Intern Med. 2018; 283: 110–20.

[55] Chowdhury PS, Chamoto K, Kumar A, Honjo T. PPAR-Induced Fatty Acid Oxidation in T Cells Increases the Number of Tumor-Reactive CD8(+) T Cells and Facilitates Anti-PD-1 Therapy. Cancer Immunol Res. 2018;10.1158/2326-6066.CIR-18-0095.

