## Supplementary material for "Altered basal lipid metabolism underlies the functional impairment of naive CD8^+^ T cells in elderly humans": Supp Mat

**Table S1. Antibodies used in flow cytometry experiments.**

| ANTIBODY | FLUOROCHROME | SUPPLIER AND REFERENCE |
| --- | --- | --- |
| Active caspase-3 | PE | BD Biosciences - 561011 |
| CCR7 | BV650 | BD Biosciences - 563407 |
| CCR7 | PE-Cy7 | BD Biosciences - 557648 |
| CD3 | BV605 | BD Biosciences - 563219 |
| CD8 | APC | BD Biosciences - 555369 |
| CD8 | APC-Cy7 | BD Biosciences - 557834 |
| CD8 | FITC | BD Biosciences - 555366 |
| CD27 | AF700 | BioLegend - 302814 |
| CD27 | BUV395 | BD Biosciences - 563815 |
| CD27 | PE | BD Biosciences - 555441 |
| CD45RA | ECD | Beckman Coulter - IM2711U |
| CD45RA | PerCP-Cy5.5 | eBioscience - 45-0458-42 |
| CD45RA | V450 | BD Biosciences - 560362 |
| CD49d | PE-Cy7 | BioLegend - 304314 |
| CD57 | Pacific Blue | BioLegend - 322316 |
| CD69 | FITC | BD Biosciences - 347823 |
| CD95 | FITC | BD Biosciences - 555673 |
| CD134 | BV711 | BD Biosciences - 563664 |
| pS6 | Pacific Blue | Cell Signaling Technology - 8520S |
| T-bet | eFluor660 | eBioscience - 50-5825-80 |

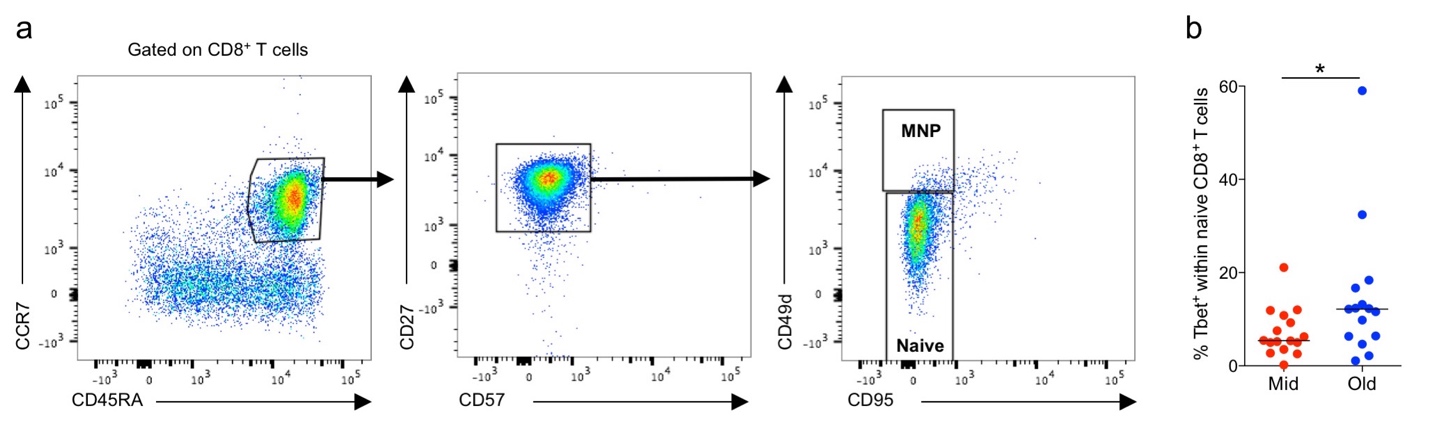

**Figure S1. T-bet expression in naive CD8^+^ T cells.** (**a**) Flow cytometric gating strategy used to exclude memory CD8^+^ T cells with a naive phenotype (MNP). (**b**) T-bet expression in naive CD8^+^ T cells (CD3^+^ CD8^+^ CD27^+^ CD45RA^+^ CCR7^+^ CD49d^−^ CD57^−^ CD95^−^) from middle-aged (Mid) and elderly individuals (Old). Each dot represents one donor. Horizontal lines indicate median values. * p < 0.05 (Mann-Whitney *U* test).

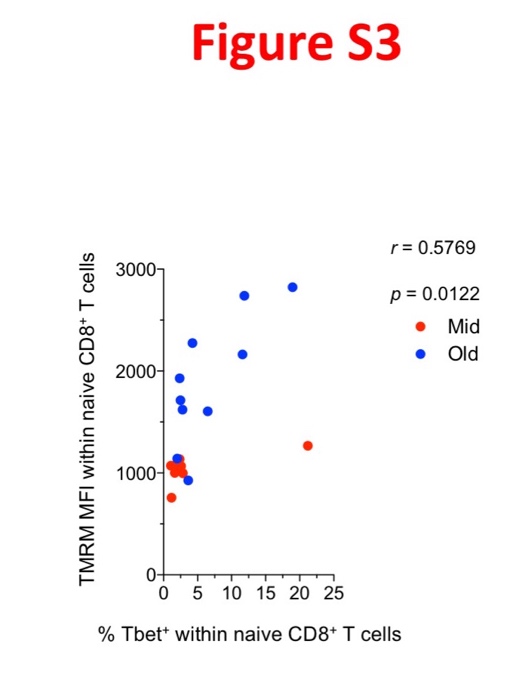

**Figure S2. Correlation between T-bet expression and mitochondrial membrane potential** **in naive CD8^+^ T cells.** Correlation between the frequency of unstimulated naive CD8^+^ T cells that expressed T-bet and basal mitochondrial membrane potential, measured by determining the mean fluorescence intensity (MFI) of TMRM. Each dot represents one donor. Significance was determined using Spearman's rank correlation.

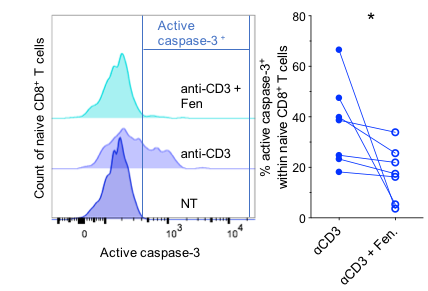

**Figure S3. Effects of fenofibrate on naive CD8^+^** **T cells.** PBMCs from elderly individuals (n = 8) were stimulated with plate-bound anti-CD3 in the absence or presence of fenofibrate (Fen). Active caspase-3 expression was measured after 24 hr. Data are shown for naive CD8^+^ T cells. Left panel: representative flow cytometry profiles. Right panel: data summary. Bars indicate mean ± SEM. * p < 0.05 (Wilcoxon signed rank test).
